# Gene therapy targeting the blood-brain barrier improves neurological symptoms in a model of genetic MCT8 deficiency

**DOI:** 10.1101/2021.12.05.471343

**Authors:** Sivaraj M. Sundaram, Adriana Arrulo Pereira, Hannes Köpke, Helge Müller-Fielitz, Meri De Angelis, Timo D. Müller, Heike Heuer, Jakob Körbelin, Markus Krohn, Jens Mittag, Ruben Nogueiras, Vincent Prevot, Markus Schwaninger

## Abstract

The solute carrier monocarboxylate transporter 8 (MCT8) transports the thyroid hormones thyroxine and tri-iodothyronine (T3) across cell membranes. *MCT8* gene deficiency, termed Allan-Herndon-Dudley syndrome, is an important cause of X-linked intellectual and motor disability. As no treatment of the neurological symptoms is available yet, we tested a gene replacement therapy in *Mct8*- and *Oatp1c1*-deficient mice as a well-established model of the disease. Here, we report that targeting brain endothelial cells for *Mct8* expression by intravenously injecting the vector AAV-BR1-*Mct8* increased T3 levels in the brain and ameliorated morphological and functional parameters associated with the disease. Importantly, the therapy resulted in a long-lasting improvement in motor coordination. Thus, the data support the concept that MCT8 mediates the transport of thyroid hormones into the brain and indicate that a readily accessible vascular target can help overcome the consequences of the severe disability associated with MCT8 deficiency.

## Introduction

The thyroid hormones (THs) thyroxine (T4) and tri-iodothyronine (T3) are critical for the function of the central nervous system (CNS). Severe hypothyroidism causes cognitive deficits and other neurological symptoms (Wood-Allum & Shaw, 2014). Mild TH deficiency has been associated with major depression and dementia (Dwyer *et al*, 2020; Tan & Vasan, 2009). In the CNS, nuclear receptors and deiodinases maintain TH homeostasis but the initial step required for the action of THs is their cellular uptake by solute carriers. Here, monocarboxylate transporter 8 (MCT8) and organic anion transporter polypeptide 1c1 (OATP1C1) transport THs *in vivo* (Bernal *et al*, 2015; Groeneweg *et al*, 2020). Mutations of the X-linked *MCT8* gene (*SLC16A2*) cause severe intellectual disability, motor dysfunction, and peripheral thyrotoxicosis (Allan-Herndon-Dudley syndrome, AHDS)(Dumitrescu *et al*, 2004; Friesema *et al*, 2004). It has been estimated that *MCT8* mutations are responsible for almost 4% of all X-linked intellectual disabilities (Visser *et al*, 2013). Affected boys appear normal at birth but show developmental delay and feeding problems in the first year of life. Some patients cannot fully control their head, speak, or walk. Other motor symptoms include muscle weakness, gait ataxia, and dystonia. TH analogues improved the peripheral thyroid status (Groeneweg *et al*, 2019; Verge *et al*, 2012), but treatment of the CNS symptoms that are key to patientś disabilities is still not available and represents an unmet medical need.

Neuropathological changes underlying symptoms of MCT8 deficiency include an altered cerebellar structure, lack of parvalbumin-positive interneurons, delayed myelination, and oligodendrocyte dysfunction (Lopez-Espindola *et al*, 2014), indicative of a global TH deficiency in the CNS. Interestingly, brain barriers, including endothelial cells, express *MCT8* into adulthood (Lopez-Espindola *et al*, 2019; Wilpert *et al*, 2020) pointing to a pivotal role of MCT8 in TH transport through the brain barriers.

Mouse studies support this hypothesis. Unlike human patients, *Mct8*^-/-^ mice do not show a neurological phenotype (Mayerl *et al*, 2014; Trajkovic *et al*, 2007); in mice but not in humans, OATP1C1 functions as a second transporter in brain endothelial cells and epithelial cells of the choroid plexus (Roberts *et al*, 2008), suggesting that OATP1C1 compensates for MCT8 deficiency. Indeed, double knockout (DKO) of *Mct8* and *Oatp1c1* (*Slco1c1*) resembles the neuropathological changes of human MCT8 mutations thus providing a suitable mouse model of the disease (Mayerl *et al*., 2014). The observation that the barrier-specific OATP1C1 can compensate for MCT8 deletion supports the assumption that MCT8 is critical for TH transport across the brain barriers (Vatine *et al*, 2017) but definitive *in vivo* evidence is lacking.

The assumption that impaired TH transport through the blood-brain barrier is the primary mechanism underlying symptoms of MCT8 deficiency suggests a novel gene therapy approach, as endothelial cells of the blood-brain barrier, unlike neural cells, are directly accessible to intravenously administered gene vectors (Körbelin *et al*, 2016). In the present study, we tested this concept in *Mct8*/*Oatp1c1* DKO mice. Our data show that AAV-mediated expression of *Mct8* in brain endothelial cells can prevent neurological pathology, identifying a critical role of MCT8 function in brain barriers and pointing to a new treatment strategy for AHDS patients normalizing TH levels in the CNS.

## Results

For targeted expression of *Mct8* in the blood-brain barrier, we generated a viral *Mct8* gene vector (Fig 1A), packaged in the AAV-BR1 capsid (AAV-BR1-*Mct8*), which selectively transduces brain endothelial cells *in vivo* (Körbelin *et al*., 2016). After incubating primary brain endothelial cells (PBECs) from *Mct8*/*Oatp1c1* DKO mice with AAV-BR1-*Mct8*, about 10% of the cells immunostained for MCT8, indicating transduction (Fig 1B). To determine whether virally transduced MCT8 was functional, we measured PBEC uptake of T3. Untreated PBECs from DKO mice did not take up T3 (Fig 1C). However, transduction of PBECs with AAV-BR1-*Mct8* enabled T3 uptake, and silychristin, an MCT8 inhibitor, blocked it, confirming that MCT8 is active after viral transduction (Fig 1C)(Johannes *et al*, 2016).

**Figure 1.**
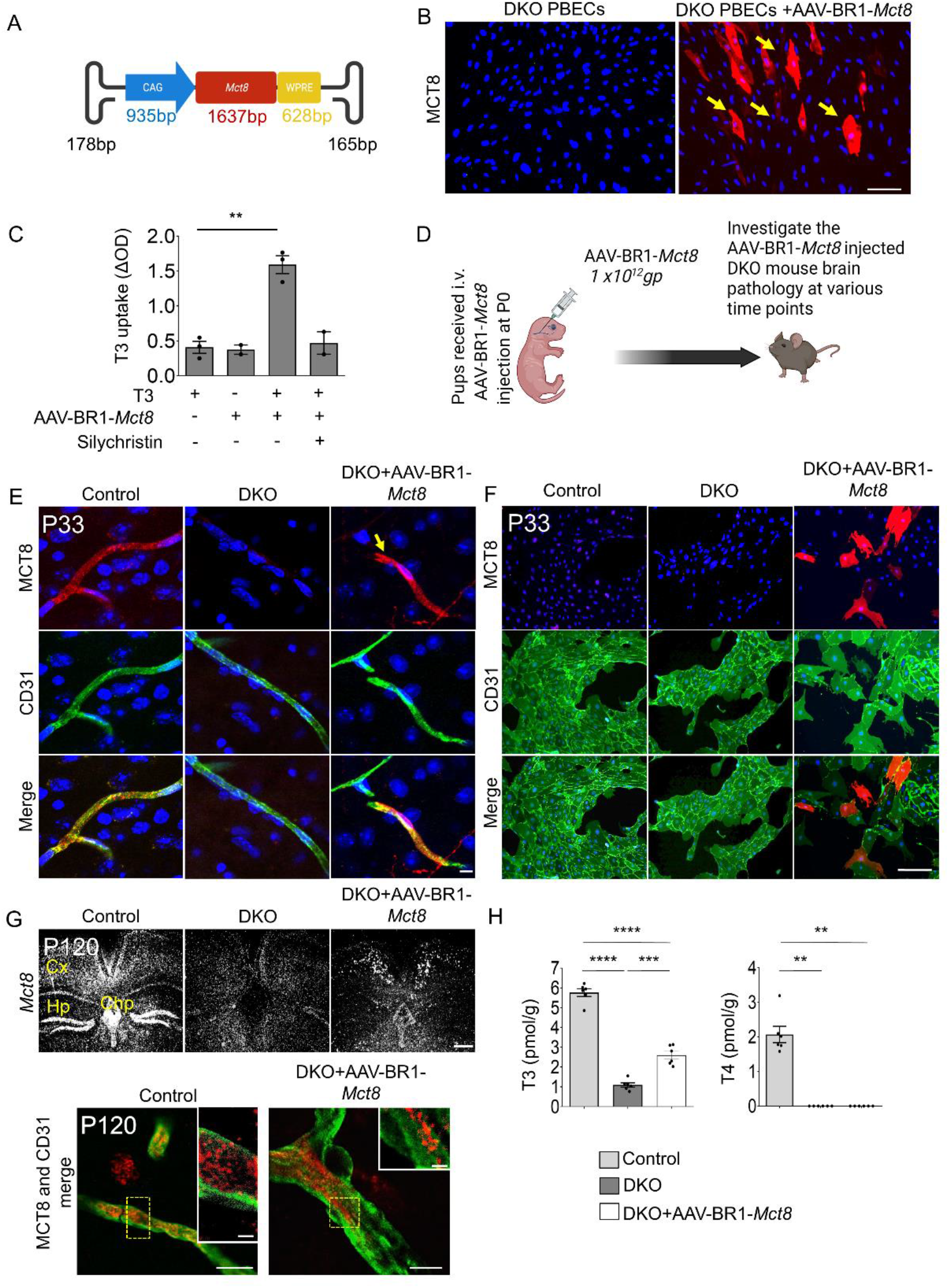
Intravenous administration of AAV-BR1-*Mct8* enables the expression of *Mct8* in brain endothelial cells and T3 transport into the CNS. (**A**) Schematic illustration of AAV-BR1-*Mct8* vectors to transduce brain endothelial cells. WPRE, woodchuck hepatitis posttranscriptional regulatory element. (**B**) After transduction of primary brain endothelial cells (PBECs) of DKO mice *in vitro* with AAV-BR1-*Mct8*, MCT8 was expressed. MCT8 was detected by fluorescence immunostaining and nuclei with DAPI. Arrows, MCT8-positive cells. Scale bar, 100 μm. (**C**) T3 uptake in DKO PBECs was enhanced by AAV-BR1-*Mct8* treatment *in vitro*. Results were obtained from three independent cell culture preparations. \*\**P*=0.0015 (unpaired t test). (**D**) Schematic of the experimental design. gp, genomic particles. (**E**) MCT8 expression (arrow) in CD31-positive endothelial cells of control or DKO mice that received AAV-BR1-*Mct8* at P0. Fluorescence immunostaining for MCT8 and CD31 was performed at P33. Scale bar, 100 μm. (**F**) MCT8 expression (arrow) in PBECs prepared from P33 mice that received AAV-BR1-*Mct8* at P0 (MCT8-positive cells, 8.6±1.1 % of CD31-positive PBECs in four independent cell culture preparations, with one mouse per cell culture preparation). (**G**) At P120, *Mct8* mRNA (upper panel) and MCT8 protein (lower panel, red) were detected in control and DKO mice treated with AAV-BR1-*Mct8* at P0. CD31, green. Scale bar, 100 μm. Cx, cortex; Chp, choroid plexus; Hp, hippocampus. (**H**) T3 and T4 concentrations in the brain of control and DKO mice that received AAV-BR1-Mct8 at P0 and were sacrificed at P21. One-way ANOVA for T3, F(2/10)=159, *P*<0.0001. One-way ANOVA for T4, F(2/10)=556.2, *P*<0.0001. Each dot represents one animal. Means ± SEM are shown. ***, *P*=0.0004; ****, *P*<0.0001 (Holm-Sidak’s posthoc test).

For *in vivo* testing, we injected AAV-BR1-*Mct8* intravenously in mice at postnatal day 0 (P0) and investigated the animals at various time points (Fig 1D). At P33, immunohistochemistry demonstrated robust MCT8 expression in brain endothelial cells and in some neurons and astrocytes, too, as previously described for the AAV-BR1 capsid (Körbelin *et al*., 2016) (Fig 1E). To quantify the transduction rate of brain endothelial cells by AAV-BR1-*Mct8*, we injected the vector in P0 mice and prepared PBECs from these animals at P33. Of PBECs derived from AAV-BR1-*Mct8*-treated mice, 8.6±1.1% expressed MCT8 (Fig 1F). However, the true transduction rate might be higher as immunofluorescence staining sensitivity for MCT8 was limited and did not detect the downregulated endogenous MCT8 levels of cultured control cells (Fig 1F)(Sabbagh & Nathans, 2020).

In the CNS, endothelial cells are non-proliferative and quiescent after mice reach an age of 14 – 30 days (Harb *et al*, 2013). Thus, loss of the non-integrating AAV vector after P33 is not expected. Accordingly, *in situ* hybridization and immunofluorescence staining showed that *Mct8* was still expressed in cerebral vessels of P120 DKO mice that received AAV-BR1-*Mct8* at P0 but not in other tissues, such as pituitary, kidney, and liver (Fig 1G, Supplemental Fig 1A, B, C), confirming the known brain tropism of AAV-BR1 after intravenous injection (Körbelin *et al*., 2016). Importantly, targeting brain endothelial cells with AAV-BR1-*Mct8* did not impair the blood-brain barrier, nor did it induce signs of endothelial cell death (Supplemental Fig 2).

As brain endothelial cell transduction sustained *Mct8* expression, we wondered whether it would affect TH levels in the brain. As shown previously, *Mct8* deletion (on an *Oatp1c1*^-/-^ background) lowered T4 and T3 concentrations in the brain (Fig 1H) (Mayerl *et al*., 2014). Importantly, AAV-BR1-*Mct8* administration to P0 mice increased cerebral concentrations of the active hormone T3 while it had no detectable effect on brain levels of the pro-hormone T4 at P21 (Fig 1H). The lack of effect on T4 levels may be due to an enhanced metabolism of T4 to T3 in DKO mice (Groeneweg *et al*., 2020). Notably, the increase in cerebral T3 levels as seen in treated DKO mice only slightly affected the hypothalamus-pituitary-thyroid (HPT) axis. *In situ* hybridization showed that *Trh* in the paraventricular nucleus and *Tshb* in the pituitary gland remained at the elevated levels characteristic for DKO mice (Supplemental Fig 3A)(Mayerl *et al*., 2014). Moreover, treatment with AAV-BR1-*Mct8* had little effect on the elevated T3 and reduced T4 serum concentrations in DKO mice (Supplemental Fig 3B) and none on the high expression of the TH-regulated genes *Gpd2* and *Dio1* in the liver or the reduced body weight characteristic of DKO mice (Supplemental Fig 4A and 4B). The fact that AAV-BR1-*Mct8* restores *Mct8* expression in the endothelial blood-brain barrier but spares tanycytes, the specialized glial cells that contain high amounts of MCT8 in the normal hypothalamus, might explain the lack of a marked effect on the HPT axis (Wilpert *et al*., 2020). Although AAV-BR1-*Mct8* would not completely normalize serum TH concentrations, in AHDS patients peripheral hyperthyroidism can be managed by the antithyroid drug propylthiouracil plus T4 or by TH analogues (Groeneweg *et al*., 2019; Verge *et al*., 2012; Wémeau *et al*, 2008).

Supply of T3 to the brain could potentially improve neurological symptoms of MCT8 deficiency. The absence of MCT8 and OATP1C1 slows down Purkinje cell dendritogenesis (Mayerl *et al*., 2014). Consequently, the molecular layer of the cerebellum consisting of dendrites of calbindin-positive Purkinje cells was thinner in untreated P12 DKO mice than in controls (Fig 2A). After injecting AAV-BR1-*Mct8*, however, the molecular layer was thicker than in untreated DKO mice (Fig 2A) and there were more interneurons expressing the marker genes GAD67 and parvalbumin in the somatosensory cortex (Fig 2A).

**Figure 2.**
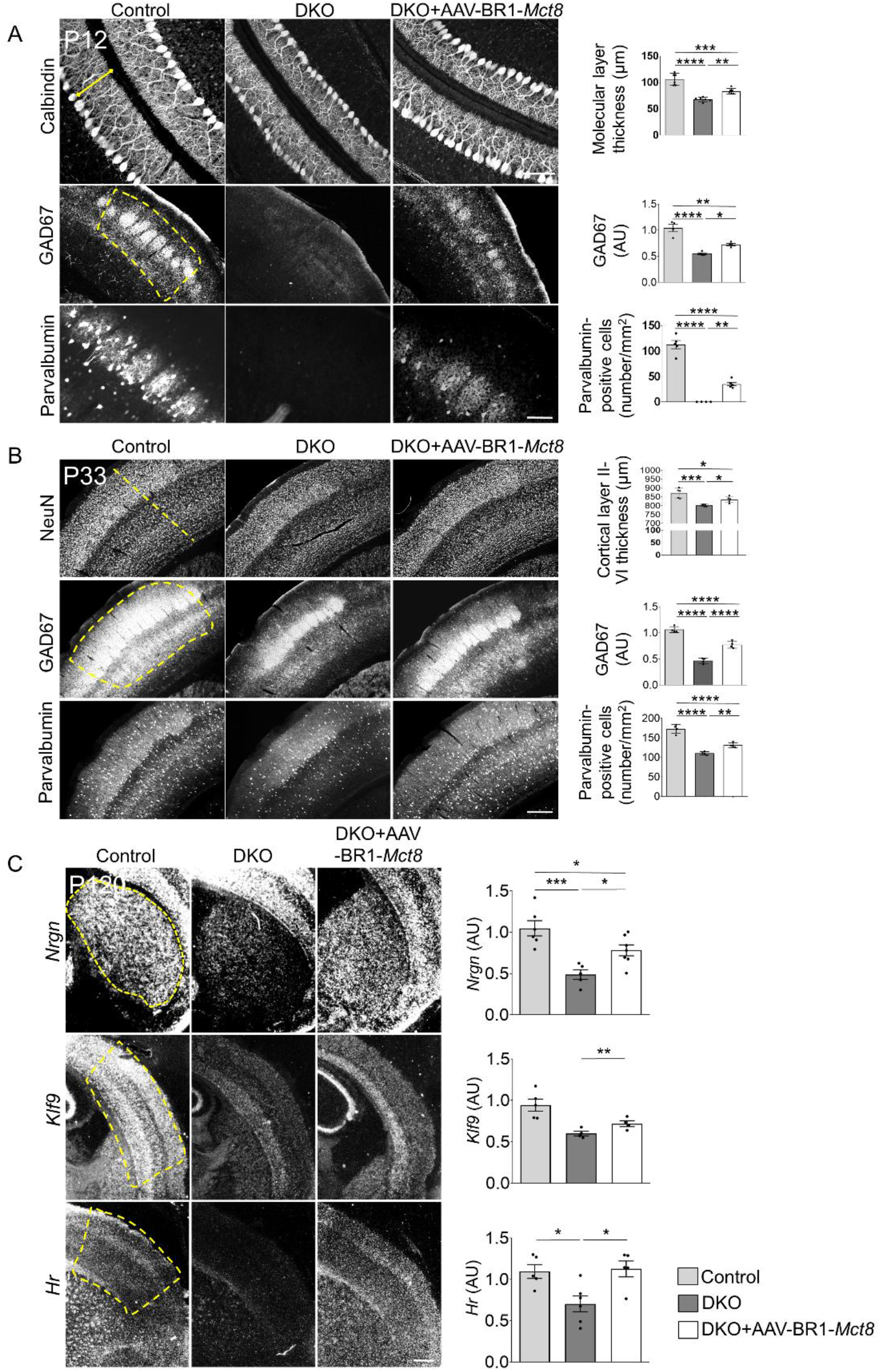
AAV-BR1-*Mct8* treatment improves neuronal morphology and gene expression. (**A**) Administration of AAV-BR1-*Mct8* at P0 to DKO mice increased thickness of the molecular layer in the cerebellar vermis as well as the relative fluorescence intensity of GAD67- and parvalbumin-positive interneurons of the sensorimotor cortex at P12. One-way ANOVA for molecular layer, F(2/15)=40.0, *P*<0.0001. One-way ANOVA for GAD67, F(2/9)=32.36, *P*<0.0001. One-way ANOVA for parvalbumin-positive neurons, F(2/11)=108.9, *P*<0.0001. Scale bar, 50 μm (upper panel) Scale bar, 200 μm (middle and lower panel) (**B**) Administration of AAV-BR1-*Mct8* at P0 improved the thickness of the somatosensory cortex layers II–VI and the relative fluorescence signal intensity of GAD67- and parvalbumin-positive interneurons in the cortex at P33. One-way ANOVA: cortical thickness, F(2/12)=16.4, *P*=0.0004; GAD67, F(2/9)=102.0, *P*<0.0001; parvalbumin, F(2/9)=63.01, *P*<0.0001. (**C**) AAV-BR1-*Mct8* injected at P0 increased the expression of *Nrgn* in the striatum and *Klf9* and *Hr* in the cortex at P120. Gene expression was investigated by *in situ* hybridization. Scale bar, 100 μm. One-way ANOVA: *Nrgn*, F(2/15)=12.8, *P*=0.0006; *Klf9*, F(2/10)=10.5, *P*=0.0035; *Hr*, F(2/13)=6.857, *P*=0.0093. Each dot represents one animal. Means ± SEM are shown. *, *P*<0.05; **, *P*<0.01; ***, *P*<0.001; ****, *P*<0.0001 (Holm-Sidak’s posthoc test).

AAV-BR1-*Mct8* treatment at P0 had a persistent effect. At P33, AAV-BR1-*Mct8*-treated DKO mice showed significantly thicker layers II–VI containing NeuN-positive neurons in the somatosensory cortex than untreated DKO animals (Fig 2B). Consistent with the findings at P12, AAV-BR1-*Mct8* treatment improved GAD67 fluorescence intensity and the number of parvalbumin-positive cells in the somatosensory cortex of DKO mice at P33 (Fig 2B).

To determine whether sustained *Mct8* expression improves neuronal function in the long term, we analyzed the expression of established TH-regulated genes by using *in situ* hybridization at P120. *Neurogranin* (*Nrgn*, Rc3) is known to be upregulated by THs in medium spiny neurons of the striatum (Iniguez *et al*, 1992). *Nrgn* expression was reduced in the striatum of untreated DKO mice (Fig 2C)(Mayerl *et al*., 2014), but after administration of AAV-BR1-*Mct8*, expression was markedly higher than in untreated DKO animals (Fig 2C). Likewise, the TH-regulated gene *Kruppel-like factor* 9 (*Klf9*) encoding a neuronal transcription factor was slightly, but nonsignificantly induced by AAV-BR1-*Mct8* treatment of DKO mice (Fig 2C). Moreover, AAV-BR1-*Mct8* injection normalized the cortical expression of the TH-regulated *Hairless* (*Hr*) in DKO mice at P120 (Potter *et al*, 2002)(Fig 2C). These findings demonstrate that AAV-BR1-*Mct8* treatment at P0 has a long-lasting effect on neuronal morphology and gene expression.

Magnetic resonance imaging and autopsy studies showed that myelination is abnormal or delayed in patients with MCT8 deficiency (Iwayama *et al*, 2021; Lopez-Espindola *et al*., 2014), reflecting the critical role of THs in postnatal myelination (Bernal *et al*., 2015). At the cellular level, THs promote differentiation of oligodendrocytes that produce myelin (Barres *et al*, 1994). As expected from the lack of THs in the CNS, DKO mice had significantly fewer Olig2-positive oligodendrocytes in the corpus callosum than controls at P33 (Fig 3A). AAV-BR1-*Mct8* treatment at P0 slightly, but nonsignificantly attenuated the decrease in Olig2-positive cells (Fig 3A). Under the influence of TH, oligodendrocytes produce myelin basic protein (MBP), a key component of CNS myelin (Farsetti *et al*, 1991). Like human patients, DKO mice present with low MBP expression in the cortex and corpus callosum (Fig 3B)(Lopez-Espindola *et al*., 2014; Mayerl *et al*., 2014). However, AAV-BR1-*Mct8* treatment at P0 increased MBP levels in the cortex of DKO mice (Fig 3B). MBP mediates myelin compaction (Readhead *et al*, 1987). Accordingly, AAV-BR1-*Mct8* treatment increased levels of compact myelin in the corpus callosum as shown by FluoroMyelin staining, indicating better gross myelination (Fig 3C).

MCT8-deficient patients suffer from coordination deficits. Like AHDS patients, DKO mice exhibited locomotor deficiencies, including impaired motor coordination and learning in comparison to littermate controls when investigated in the rotarod test at ∼ P120 (Fig 4A)(Mayerl *et al*., 2014).

**Figure 3.**
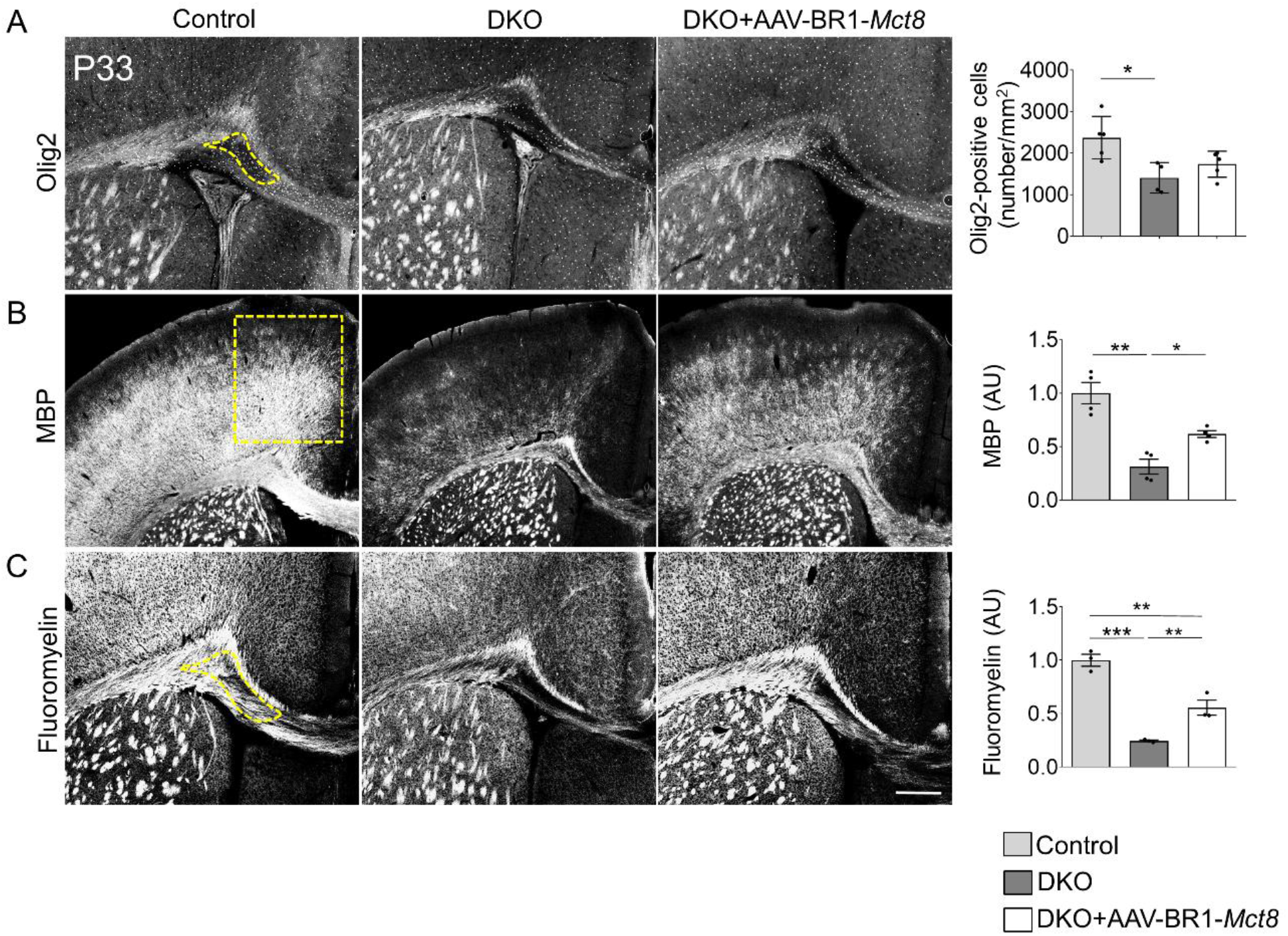
AAV-BR1-*Mct8* treatment increases oligodendrocyte numbers and myelination. Treatment of DKO mice at P0 improved several parameters of myelination at P33. (**A**) The reduced number of Olig2-immunopositive oligodendrocytes in the corpus callosum was slightly but nonsignificantly increased in DKO treated with AAV-BR1-*Mct8*. Scale bar, 100 μm One-way ANOVA, F(2/11)=6.595, *P*=0.0131. *, *P*<0.05 (Holm-Sidak’s posthoc test). (**B**) AAV-BR1-*Mct8* improved myelin basic protein (MBP) levels in the cortex of DKO mice. MBP was detected by immunostaining. Scale bar, 100 μm. Welch’s ANOVA test, W (2/4.96)=14.8, *P*=0.0081. *, *P*<0.05; **, *P*<0.001 (Tamhane’s T2 posthoc test). (**C**) Compact myelin stained by Fluoromyelin was diminished in the cortex of untreated DKO but improved by treatment with AAV-BR1-*Mct8*. One-way ANOVA, F(2/6)=53.15, *P*=0.0002. **, *P*<0.01; ***, *P*<0.001 (Holm-Sidak’s posthoc test). Each dot represents one animal. Means ± SEM are shown.

**Figure 4.**
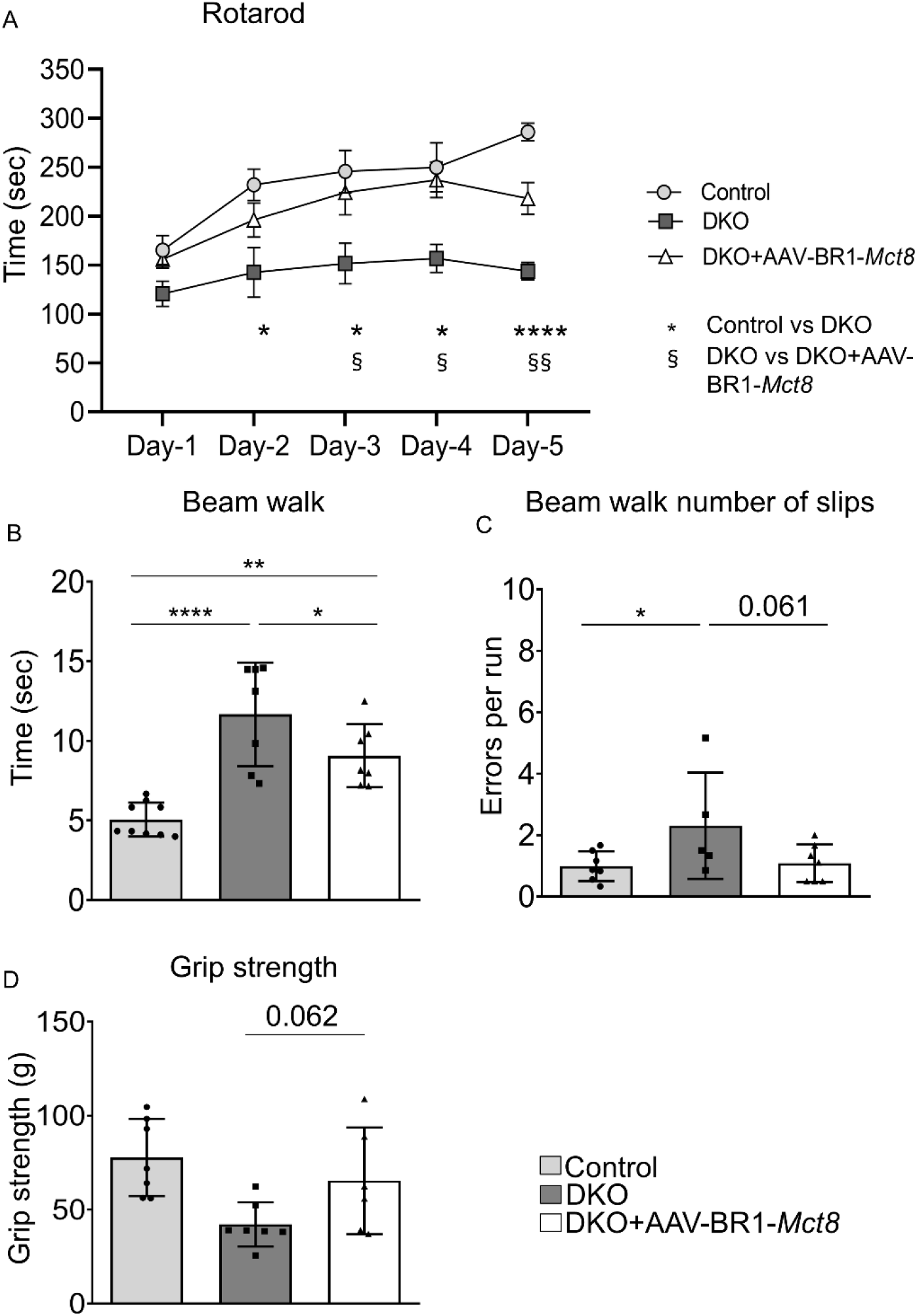
AAV-BR1-*Mct8* treatment improves motor function in DKO mice. (**A**) AAV-BR1-*Mct8* treatment at P0 prolonged the time DKO mice were able to balance on the rotarod as a sign of improved coordination and motor learning. Mice were assessed on 5 consecutive days starting at P120. Repeated-measures ANOVA, F(2/14)=11.85, *P*=0.001. (**B, C**) AAV-BR1-*Mct8* treatment reduced the time DKO mice needed to cross the beam and led to a trend towards fewer errors, indicating better motor coordination. One-way ANOVA for time, F(2/20)=18.56, *P*<0.0001. Kruskal-Wallis test for errors, *P*=0.215. (**D**) AAV-BR1-*Mct8* treatment increased the grip strength of DKO mice in comparison to control animals. One-way ANOVA, F(2/17)=5.25, *P*=0.017. *,§*P*<0.05, **,§*P*<0.01, \*\*\*\**P*<0.0001 (Holm-Sidak’s posthoc test). ns, nonsignificant. Each dot represents one animal. Means±SEM are shown.

However, a single intravenous injection of AAV-BR1-*Mct8* at P0 significantly improved DKO mice performance and motor learning (Fig 4A). In the beam walk, another test of coordination and balance, DKO mice treated with AAV-BR1-*Mct8* crossed the beam faster than untreated DKO and tended to make fewer errors (Fig 4B, C). Moreover, grip strength tended to be greater in AAV-BR1-*Mct8-*treated DKO mice (Fig 4D).

## Discussion

Our study reports a novel gene therapy for MCT8 deficiency, a major cause of X-linked intellectual disability. A previous study expressed *MCT8* in mice with the help of the AAV9 capsid that penetrates the blood-brain barrier and transduces neural cells but expression in parenchymal cells did not correct low TH concentrations in the brain after intracerebroventricular injection of the vector, presumably because the blood-brain barrier transport of THs was still compromised (Iwayama *et al*, 2016). Moreover, the unspecific tropism of AAV9 and its high liver transduction are associated with side effects (Chand *et al*, 2021; Walia *et al*, 2015). In contrast, targeting the small endothelial cell population in the present study was sufficient to supply T3 to the brain and to ameliorate symptoms of MCT8 deficiency in a mouse model, although the transduction rate of endothelial cells was rather low. Thus, endothelial cells are a critical site for MCT8 function after birth.

Normal brain development requires TH action already before birth, but how THs cross the prenatal blood-brain barrier is still unclear. Human brain vessels express MCT8 already at gestational week 14 (Lopez-Espindola *et al*., 2019). Lower brain TH concentrations in an MCT8-deficient fetus suggest that MCT8 plays a role in fetal TH transport (Lopez-Espindola *et al*., 2014). On the other hand, patients with MCT8 deficiency are normal at birth and only develop symptoms during the first year of life (Dumitrescu *et al*., 2004; Friesema *et al*., 2004; Groeneweg *et al*., 2020). Our study now provides functional data that postnatal reconstitution of MCT8 is sufficient to prevent the neurological phenotype of the disease, at least partially.

MCT8 expression could be directed to the neonatal blood-brain barrier by using an AAV vector with high selectivity for brain endothelial cells (Körbelin *et al*., 2016). This approach improved transcriptional and morphological parameters of the disease already after 12 days. Importantly, the effect persisted at the behavioral and transcriptional level into adulthood. Thus, brain endothelial cell-specific gene therapy emerges as a novel treatment of the disabling neurological signs of AHDS. The intravenous route and postnatal administration may facilitate translation to clinical application.

## Materials and methods Mice

Breeding pairs were set up among Mct8^-/+^;Oatp1c1^-/+^ and Mct8^-/y^;Oatp1c1^-/+^ mice to obtain Mct8^-/-^;Oatp1c1^-/-^ or *Mct8*^-/y^;*Oatp1c1*^-/-^ (DKO) and Mct8^-/+^;Oatp1c1^-/+^ or Mct8^+/y^;Oatp1c1^-/+^ control mice, as reported previously (Mayerl *et al*., 2014). Detailed information is provided in Supplemental Methods. Mice were randomized into treatment groups. Experimenters were blinded to group allocation in the phenotype analysis.

### Statistical analysis

*P* values ≤ 0.05 were considered statistically significant. We did not exclude outliers, unless indicated. ANOVA and t-test were only applied if assumptions were met, i.e., datasets were examined for Gaussian distribution using the D’Agostino-Pearson or Kolmogrov-Smirnov test, aided by visual inspection of the data, and homogeneity of variances by Brown-Forsythe test. Mostly, Holm-Sidak posthoc analysis was applied to test the significance between groups.

### Study approval

All animal experiments were performed according to German animal welfare regulations, and experimental protocols were approved by the local animal ethics committee (No. 11-1-17, MELUND, Kiel, Germany).

## Author contributions

J.M. and M.S. designed the study. S.M.S., A.A.P., H.K., H.M.F., M.D.A., J.K., M.K., and C.L.S. performed experiments and analyzed the data. H.H., T.D.M., R.N., V.P., and J.M. provided tools, samples, and conceptual support. S.M.S, A.A.P., R.N., V.P., and M.S. drafted the manuscript. All authors revised the manuscript for important intellectual content.

## Acknowledgement

We thank Ines Stölting, Wiebke Brandt, Beate Lembrich, Frauke Spiecker (Pharmacology), and Christian L. Schmidt (Isotope laboratory) for their kind support. This work was supported by the European Research Council (ERC) Synergy Grant-2019-WATCH-810331 to V. P., R. N., and M. S., by grants of the Deutsche Forschungsgemeinschaft (MU-3743/1-1 to H.M.F., CRC/TR296 to H.M.F., T.M., H.H., J.M. and M. S.) and by the Sherman family funds to H.H..

